# Predicting Protein Crystal Solvent Content from Patterson Maps Using Machine Learning

**DOI:** 10.1101/2025.09.24.678396

**Authors:** David McDonagh, Daniel J. Rigden, David G. Waterman, Ronan M. Keegan

## Abstract

Estimating the solvent content of protein crystals is fundamental to identifying the correct symmetry and phasing of the unit cell. Typically, the number of molecules in the asymmetric unit is not known and probabilistic methods are used based on statistics derived from the Protein Data Bank (PDB). These methods tend to predict the number of molecules incorrectly in around 20% of cases, which can significantly impede the structure solution pipeline. Here multiple machine learning approaches are investigated to predict solvent content using Patterson Maps. Several architectures are shown to give a significant improvement over current approaches, with prediction errors being reduced by over 50%. In addition, the potential of embedded representations of Patterson Maps for clustering is demonstrated, which could lead to new approaches for identifying similar structures when processing novel data.

## 1 Introduction

A key step in the structure determination process of protein crystals is an estimate of the molecular unit cell content. This can be performed after diffraction data has been indexed with a candidate lattice using modern diffraction reduction software [1–3]. Identifying the potential number of molecules in the unit cell, and the corresponding solvent content, plays a key role in identifying the overall symmetry of the cell, in addition to addressing the phase problem required to calculate the electron density [4]. More than five decades ago Matthews analysed the solvent content of 116 crystals of globular proteins, finding a range of 27-78%, with the most common value being 43% [5, 6]. Assuming a constant specific protein density, the relationship between solvent content and the number of molecules in the asymmetric unit was quantified in terms of a coefficient, *V*_*M*_

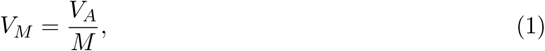

where *V*_*A*_ is the crystal asymmetric unit volume and *M* is the molecular weight, which relates to the solvent content [7], *V*_*S*_, as

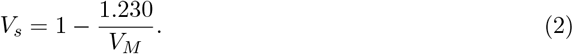

Hence if the unit cell and number of molecules in the asymmetric unit are known, the solvent content can be calculated. In the structure solution pipeline this is generally not the case, however, and the number of molecules must be estimated. One approach to this has been to incorporate the Matthews Coefficient (1) into a probabilistic estimator, taking into account the correlation observed in the PDB between solvent content and resolution [8]. This is given by

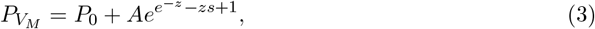

where

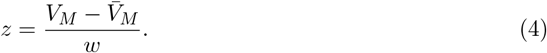

Here *P*_0_, *A, s*, and *w* are empirical parameters optimised over 12 resolution ranges from a subset of the PDB [8], and 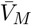 is the average value of the Matthews Coefficient for a given resolution range. Such an approach can be used to estimate the likelihood of different numbers of molecules in the asymmetric unit and hence different solvent content values. This has been implemented in packages such as MATTHEWS_COEF in CCP4 [9] for more than two decades, and is now routinely used as part of the structure solution pipeline.

In this study the performance of MATTHEWS_COEF was investigated using a set of 80,691 protein crystals from the PDB [10], released prior to July 2022, where structure factor amplitudes or intensities had been deposited. Entries that contained only protein molecules were included to enable uniformity and avoid any differences that might arise when RNA and/or DNA molecules are present in the crystal. Some redundancy in the selection from the PDB was tolerated in order to accumulate as large as possible a dataset for the training. Reported solvent content values were compared with a recalculated solvent content. Where these values differed, the case was examined in detail to ascertain the correct solvent content or, in rare cases, eliminate the example from the dataset due to some peculiarity of the deposited data. For the purposes of this study solvent content was selected as the target for the calculation rather than molecular content. A key reason for this is estimating solvent content makes no assumption about the molecular content of the crystal such as atom types, sequence, and molecule type. This helps eliminate any potential bias due to an assumed cell content when considering designing new approaches for prediction.

Fig. 1a shows how MATTHEWS_COEF incorrectly predicts the number of molecules in the asymmetric unit around 20% of the time. Prediction error is also found to increase with the number of molecules, where beyond around 20 molecules in the asymmetric unit predictions are almost always incorrect (Fig. 1b). Poorly predicted solvent content values can be computationally expensive in the structure solution pipeline. This is one of the critical parameters that determines success in phasing by molecular replacement, for example.

**Figure 1.**
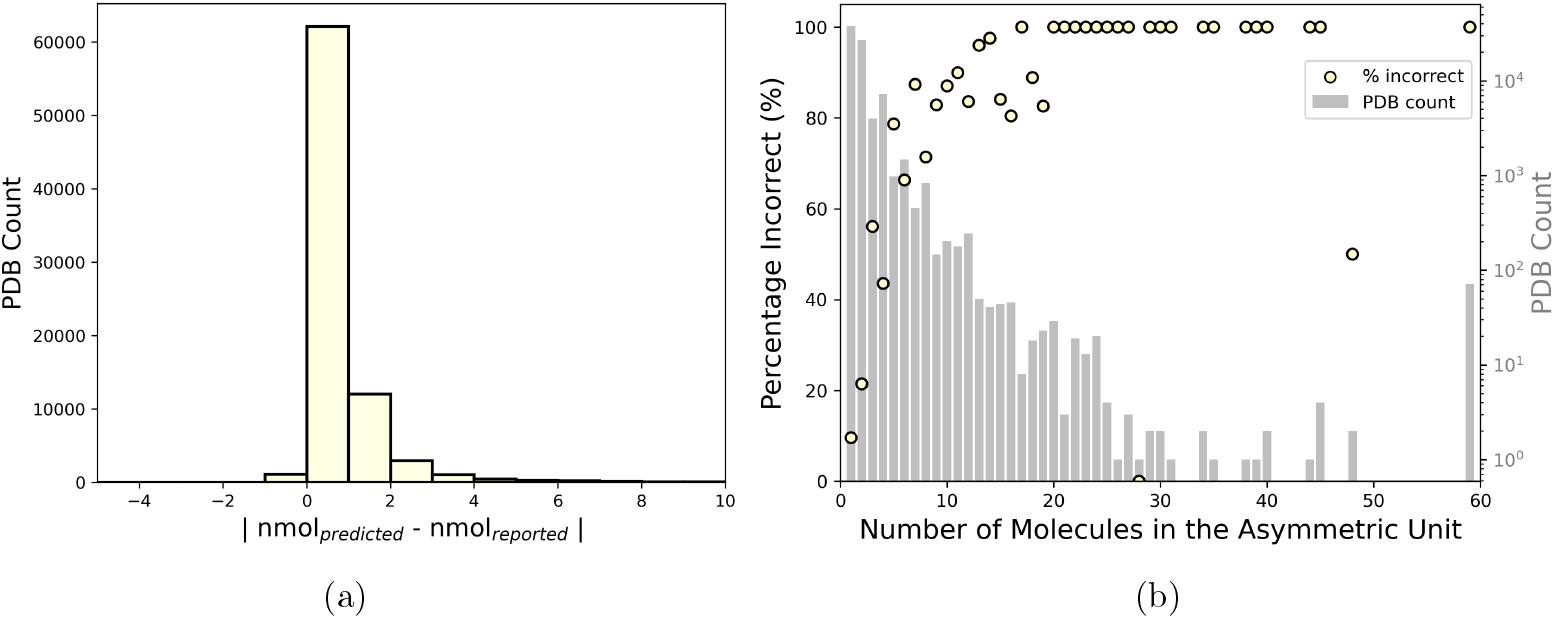
A comparison of the number of molecules predicted using MATTHEWS_COEF versus that reported in the PDB for 80,691 structures. (a) A histogram of the differences between predicted and reported values. (b) How the prediction error changes with number of molecules in the asymmetric unit. For MATTHEWS_COEF predictions, molecular weights were calculated from SEQRES records using the Biopython package [11], and the highest probability prediction was selected in each case.

In recent years, machine learning methods have increasingly been used in crystallography [12, 13], including the CCP4 suite [14–16]. These data-driven approaches are ideal for protein crystallography, owing to the now over 200,000 experimental structures available in the PDB. Supervised learning methods attempt to map a relationship between data representations and properties using highly flexible functions, consisting of architectures that are adapted to learning layered relationships (e.g. neural networks). The success of their application can allow the exploitation of complex representations of protein crystals that are difficult for researchers to interpret. One example of this is the Patterson map, which is defined for every position in the unit cell as

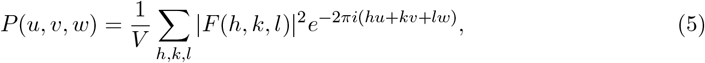

where *V* is the cell volume, and *F|*(*h, k, l*)*|* is the structure factor amplitude at Miller index (*h, k, l*), which is obtained from the square root of the measured intensity from the corresponding diffraction pattern reflection [17]. Although these maps contain no phase information, peaks in the distribution relate to interatomic distances within the unit cell, where the amplitude of the origin peak is proportional the number of atoms. The Patterson map should, therefore, contain significant information about the solvent content. However, this is difficult to interpret directly due to peak overlap and the map containing *n*^2^ peaks for *n* atoms in the unit cell.

Given these maps are essentially three-dimensional images, they are well suited to established computer vision machine learning architectures, such as Convolutional Neural Networks (CNNs) [18], and Vision Transformer models (ViTs) [19]. CNNs have previously been used to map Patterson maps directly to atom coordinates using synthetic datasets of maps with the same dimensions [20]. In this case a custom network of 12 3D convolutional layers was trained on maps from randomly generated positions of 10 atoms. More recent work followed a similar approach, using the CNN U-Net architecture [21] to predict electron density maps from Patterson maps, based on a synthetic dataset derived from structures in the PDB [22].

In this work the aim is not to predict the electron density itself, but instead focus on the more tractable mapping of predicting the solvent content from Patterson maps. The goal is not to replace phasing methods, but rather to significantly improve accuracy and efficiency in the phasing pipeline. This work is also distinct in using a large dataset of experimental structures directly from the PDB, with varied unit cell dimensions.

## 2 Methodology

The dataset described in the previous section was used throughout this work. Patterson maps were generated using the FFT software in CCP4, using ECALC to obtain normalised amplitudes as

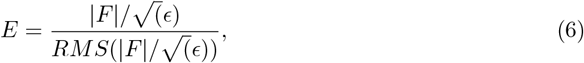

where *|F|* is the structure amplitude, *ϵ* is a symmetry factor increasing mean intensities corresponding to the Laue group symmetry, and *RMS* is a function of resolution, normalising the data such that *< E*^2^ *>*= 1. Ground truth solvent content values were taken from the PDB, with all machine learning model predictions compared against solvent content values calculated using the MATTHEWS_COEF package in CCP4 [9]. The distribution of all unit cell lengths for the dataset is shown in Fig. 2. As with general computer vision tasks, the different unit cell sizes (and hence Patterson maps) can be accounted for via scaling, padding, cropping, or global pooling layers within a network architecture. In this work, models trained on the maps themselves appeared to learn best when using a consistent Cartesian grid of size (256, 256, 256), where Patterson maps were interpolated at values of 0.5 °*A* using the Gemmi software package [23]. This size was selected based on looking at the average unit cell lengths in the dataset and rounding to the nearest power of two (i.e 128) to make them more amenable to neural network architectures.

**Figure 2.**
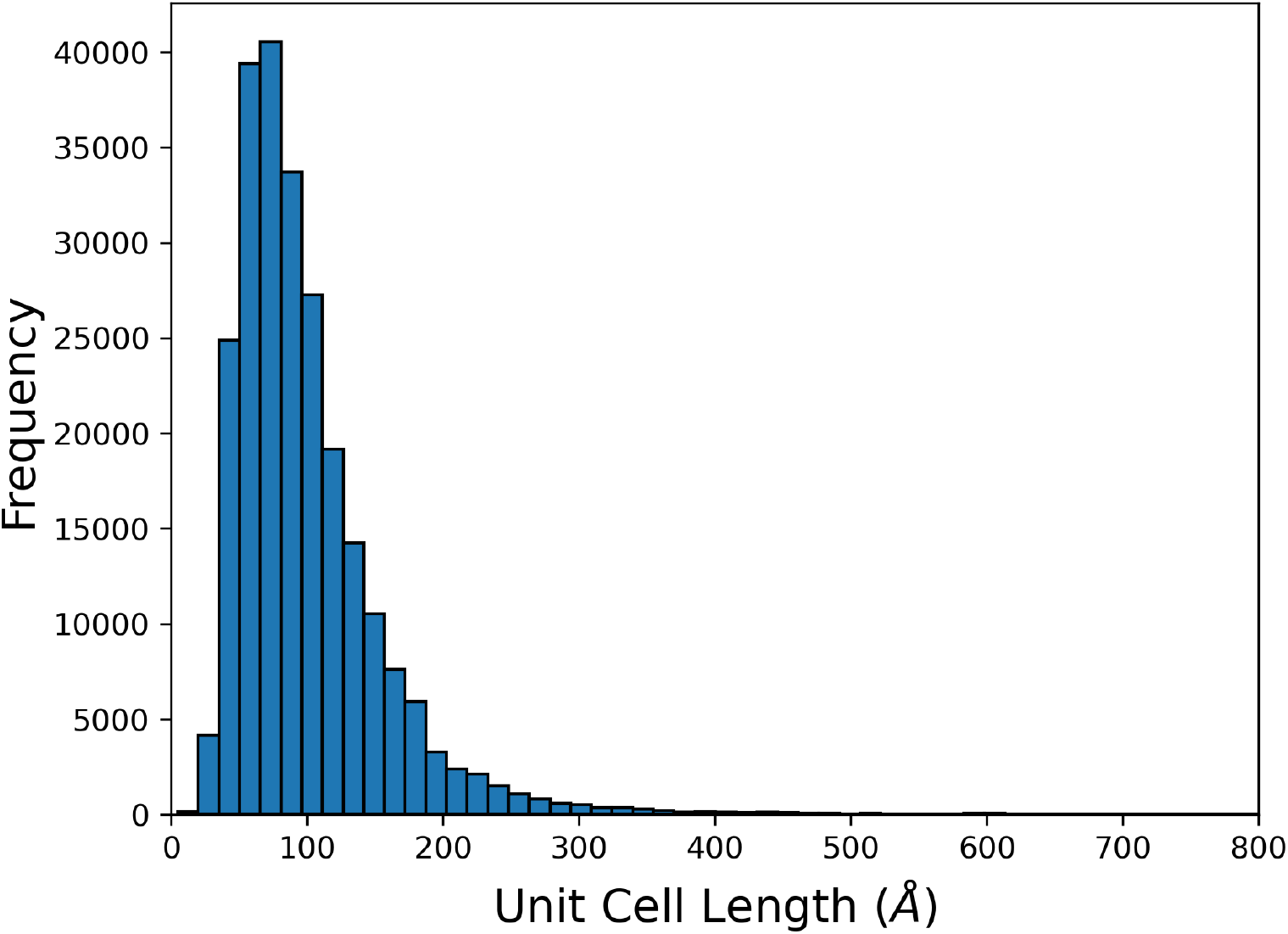
The distribution of unit cell lengths across the dataset of 80,691 protein crystal structures taken from the PDB. The min, max, median and mean values are 4.7, 1526.4, 85.7 and 99.2 °A, respectively.

The distribution of reported solvent content for the dataset is given in Fig 3. The dataset was randomly split into training, validation, and test sets, using 75%, 5% and 20%, respectively. Given the non-uniform distribution of solvent content values, the data was first binned by increments of 10%. Random splitting was then done per bin to better ensure each set accurately represented the overall distribution.

**Figure 3.**
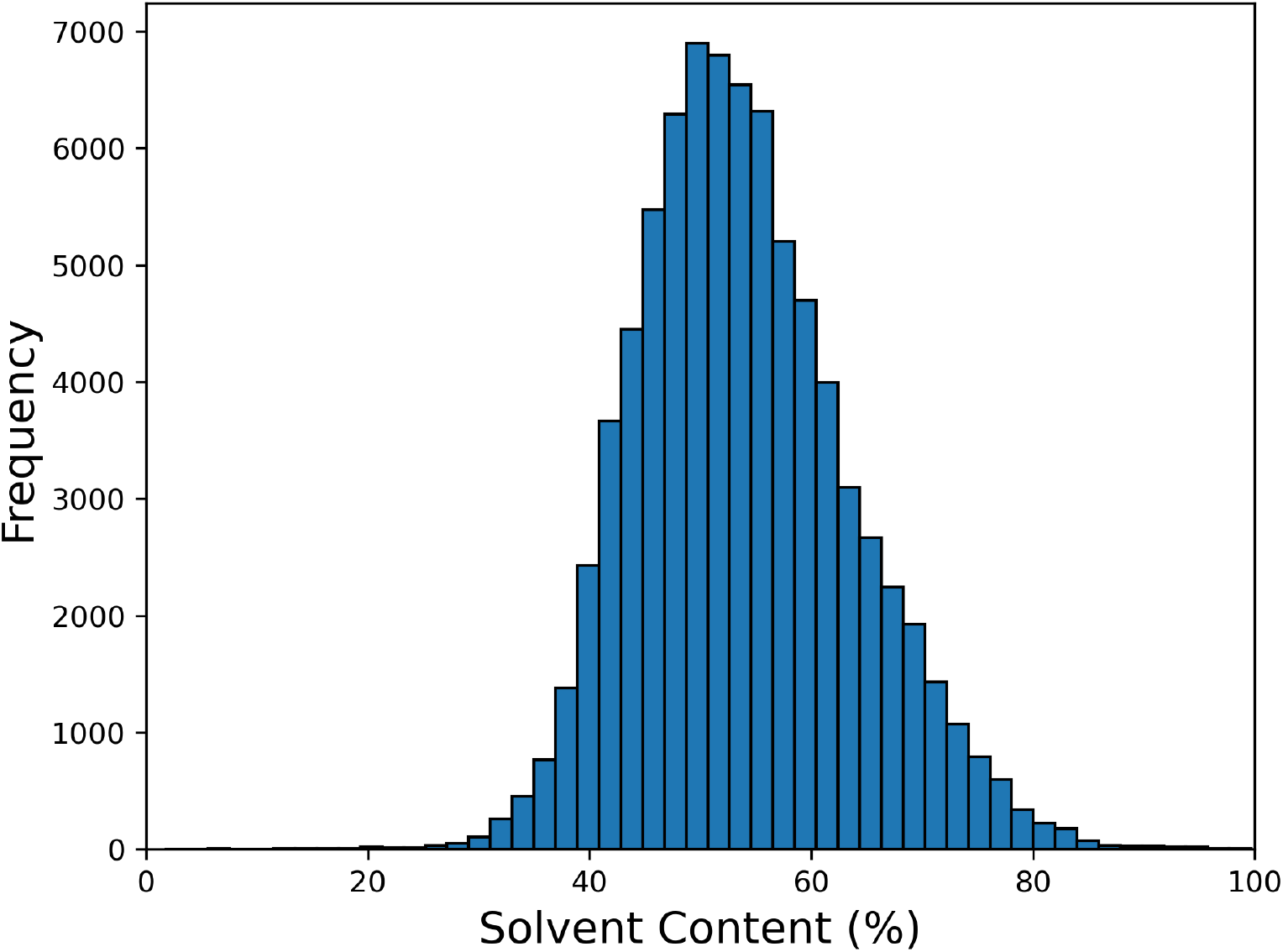
The distribution reported solvent content values for the dataset used in this work.

To map protein crystal data to solvent content three different approaches were taken: manual feature selection of Patterson map and crystal properties (Section 2.1), and Patterson maps used directly with adapted versions of the CNN U-Net and ViT-V-Net architectures [21, 24] (Sections 2.2 and 2.3, respectively). In each case the aim was to minimise the mean squared error (MSE)

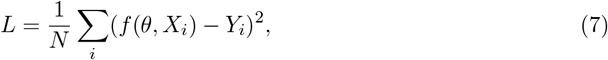

where *Y*_*i*_ is the reported solvent content for the *ith* training set structure, and *f* (*θ, X*_*i*_) is the machine learning model, defined as a function of both the Patterson map and crystal properties *X*_*i*_, and learnable parameters *θ*. In light of the unbalanced distribution of solvent values given in Fig. 3, for the two neural network models the cost function was augmented with a regularisation term,

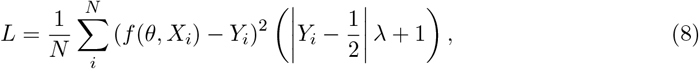

with coefficient *λ*. This then weights the contribution to the cost function from samples at the tail ends of the distribution more heavily, with the aim of further minimising overfitting to predictions towards the mean of the training set distribution. In this work a *λ* value of 8 was used, based on initial performance in predicting training set structures with solvent content values at the extremes of the dataset. This augmented cost function was not used with the manual selection model due to restraints within the used machine learning package and the model described in Section 2.1 being less susceptible to overfitting.

### 2.1 Manual Feature Selection

Previous work has highlighted resolution as the dominant property correlated with solvent content, with other properties such as molecular weight only showing a weak correlation [7]. In this work a number of descriptors were examined using the training set. A summary is given in Table 1, along with correlation coefficients with the reported solvent content. Correlation plots are given in Appendix A. The descriptors investigated were chosen based on assumptions about what features in the Patterson map might be important with respect to solvent content. Higher standard deviation of the Patterson map, for example, was assumed to be more likely for maps with low solvent content as a greater number of interatomic distances should be possible. Similarly, heavily solvated structures were assumed to have a higher peak to noise ratio, as the possible interatomic positions would be reduced.

**Table 1:** Features used in this work for the Random Forest model and their correlation to the reported solvent content for the training set.

| Feature | Correlation | Description |
| --- | --- | --- |
| mean grad | -0.693 | The mean of the magnitude of the gradient of the Patterson map, calculated using the Sobel kernel [28] as a 3D convolution. |
| std | -0.688 | The standard deviation of the Patterson map. |
| per grad | -0.683 | The 90th percentile of the mean of the magnitude of the gradient of the Patterson map, calculated using the Sobel kernel [28] as a 3D convolution. The percentile used was optimised with the training set to maximise correlation with solvent content. |
| volume | 0.682 | The log of the volume of the unit cell. |
| low density | 0.664 | The proportion of values in the Patterson map below a given threshold, normalised with respect to the size of the map, optimised using the training set to maximise correlation with solvent content. |
| resolution | 0.546 | The observed resolution of the structure. |
| max grad | -0.541 | The max of the magnitude of the gradient of the Patterson map, calculated using the Sobel Kernel [28] as a 3D convolution. |
| mw | 0.447 | Molecular weight. |
| variance | 0.281 | The variance of Patterson map values as a histogram, with bin width size optimised using the training set to maximise correlation with solvent content. |
| peak noise ratio | -0.272 | An estimate of the peaks in the Patterson map relative to noise. Peaks were taken as values above $\mu + 2\sigma$. The ratio was then calculated as $\frac{\mu_{peaks} - \mu}{\sigma}$. |
| kurtosis | 0.263 | A measure of the distribution of values in the Patterson map defined as $\frac{\mu^4}{\sigma^4}$, where $\mu$ is the mean and $\sigma$ is the standard deviation. |
| skew | 0.235 | A measure of the skew of values in the Patterson map. |

**Table 2:** The energy cost and associated carbon footprint to train each of the models. Values for the Random Forest model were obtained using CodeCarbon on both descriptor generation and training (all done on the CPU) [36]. These results were then compared against those obtained from the Green Algorithms calculator [37] and found in broad agreement. Values for ViT-Solv and U-Solv were obtained using the Green Algorithms calculator [37].

| Model | Training time (hours) | Energy Use (kWh) | Carbon Footprint (kgCO<sub>2</sub>e) |
| --- | --- | --- | --- |
| Random Forest | 27 | 3.24 | 0.36 |
| ViT-Solv | 174 | 170.53 | 18.96 |
| U-Solv | 165 | 161.71 | 17.98 |

As these features are all scalar values, they can be used in the plethora of machine learning models available that learn from a vector input. To identify a suitable model, the Tree-based Pipeline Optimization Tool (TPOT) was run for 2000 generations [25]. Here a number pipelines consisting of different machine learning models from the Python Scikit-learn package [26] were analysed using 5-fold cross validation. This identified a random forest model [27] after the data has been appended with components obtained from principal component analysis. Specifically, the manual feature model pipeline consists of the following steps:

#### Manual Feature Selection Model Generation

1. Centre each feature in the training set and calculate the covariance matrix
2. Calculate principal components
3. Append these principal components to the original descriptors as input and the solvent content as output
4. Fit an ensemble of 100 decision trees with a squared error loss function, each using a bootstrap dataset with a random subset of features
5. Take the average of all decision trees to give the overall solvent content prediction on the test set

For the remainder of this work this model is referred to as the Random Forest model.

### 2.2 CNN Architecture

In this study the U-Net CNN architecture was used [21], owing to its versatility in extracting data from 2D and 3D images [29], as well as related areas in structural biology [14]. The version used in this study was the model adapted to 3D data, enhanced with residual blocks and spatial squeeze and channel excitation blocks [30–32]. The basic 3D U-Net architecture consists of a series of 3D convolutions and pool layers to downsample 3D feature vectors (the encoder). A decoder component then upsamples and combines encoder features via skip connections [29]. For the purposes of this study, only the decoder element was used as the aim was not to reproduce an image of the same dimensions as the input but instead a scalar value (the solvent content). The overall architecture was therefore adapted to add a final fully connected layer to the last pooling layer of the encoder, with the result passed through a sigmoid function to give a value between zero and one. The full model details are given in Appendix B. To avoid confusion with the standard U-Net architecture, for the remainder of this work this model is referred to as U-Solv.

### 2.3 Transformer Model Architecture

Vision transformer models were inspired by models first introduced in natural language processing [19, 33]. Rather than extracting hierarchical features via convolutions with the image pixel data, images are divided into a series of patches, analogous to words. These patches are then flattened and linearly projected into a lower dimensional embedding before being processed with self attention and feed forward layers. In this work a 3D adaption of the ViT-V-Net model was used [24, 34]. This model consists of an initial 3D CNN to extract features, combined with a transformer model to divide the feature maps into patches to learn longer range, correlated features. As with the U-Solv model only the encoder component was used. The final layer of patches was then averaged and fed into to a fully connected layer, again summing to a single output mapped with a sigmoid function to give a value between zero and one. The full model details are given in Appendix C. Again to avoid confusion with the standard architecture, for the remainder of this work this model is referred to as ViT-Solv.

### 2.4 Network Training

Both neural networks were trained on the training set only, using a batch size of 8 (limited by computational hardware), a learning rate of 1*e*^−4^, the modified cost function given in eq (8), and the Adam optimisation algorithm [35]. For the ViT-Solv model, a dropout probability of 0.1 was applied to the sum of patch embeddings and position embeddings, and to the multilayer perceptrons (MLPs) inside each transformer block. After initially training for 15 epochs, training was stopped as soon as the validation error diverged from the training error. For the U-Solv model this was at epoch 15, whereas for the ViT-Solv model this occurred at epoch 29.

## 4 Results and Discussion

All four models were tested using the test set of 16,094 structures that were not used in any part of model training. The overall performance for each of the models is shown in figures 4 and 5. All machine learning models are observed to perform well with root mean squared errors (RMSEs) approximately half of that obtained with MATTHEWS_COEF (Fig. 4a). The number of structures in the test set with predicted values within 10% of the reported value is found to improve from 77.5% using MATTHEWS_COEF to 96.38% using the U-Solv model.

**Figure 4.**
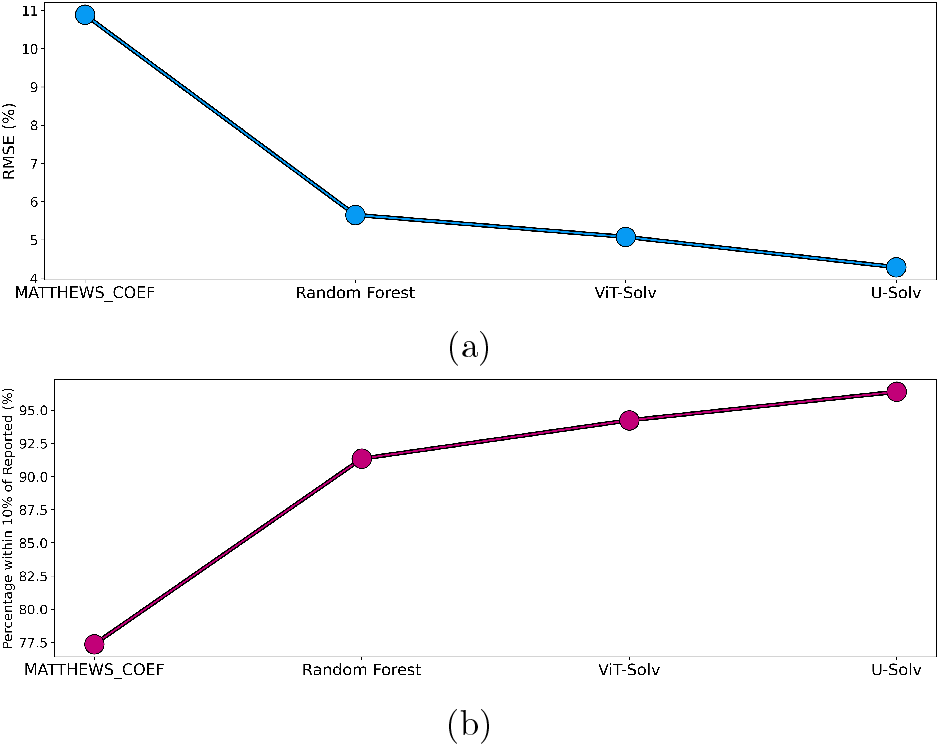
The performance of each of the four models in terms of (a) root mean squared error (RMSE) and (b) the number of structures with predicted solvent content values within 10% of the reported values.

**Figure 5.**
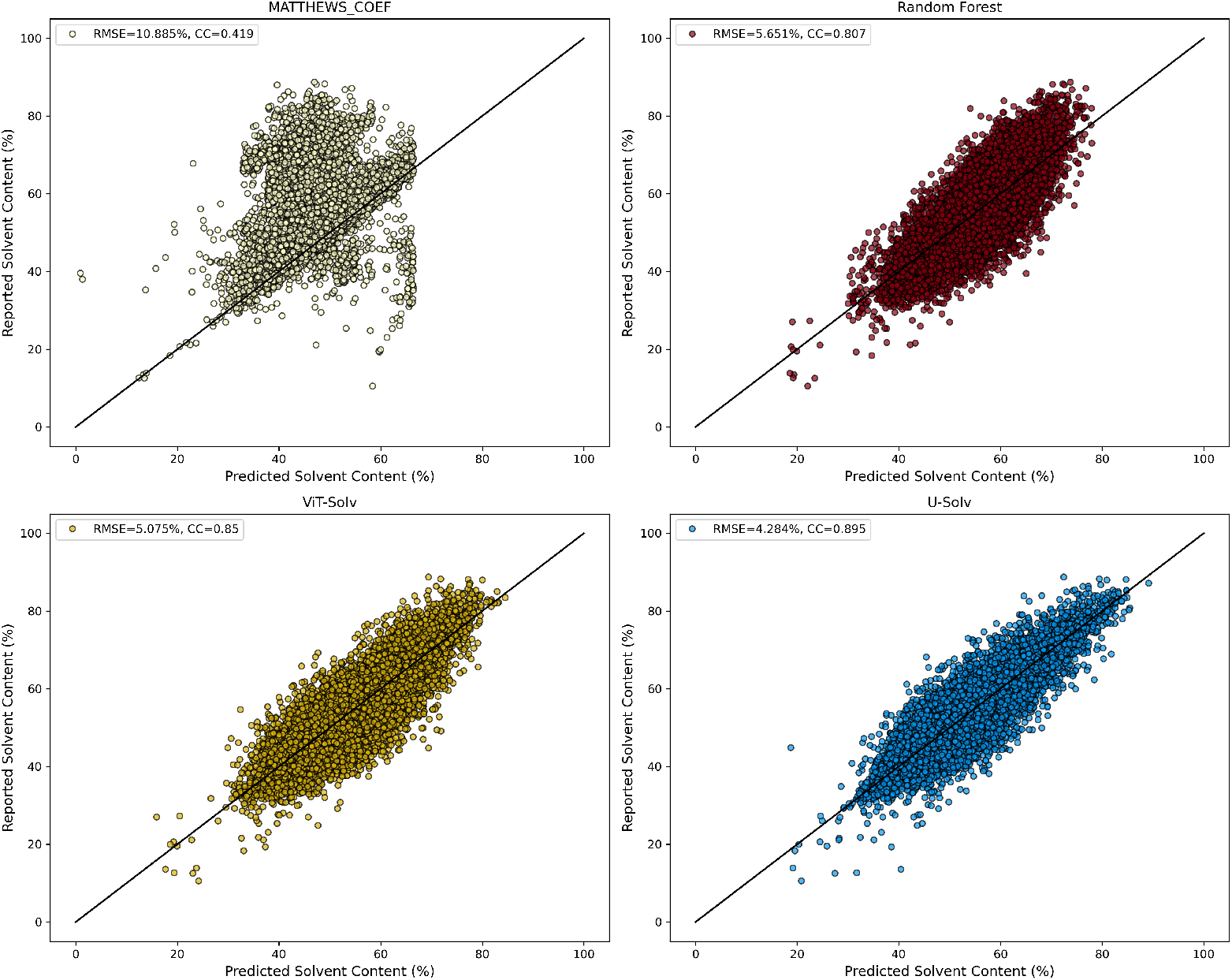
Correlation plots for predictions versus reported solvent content values for the structures in the test set used in this study. As in Fig. 1, for MATTHEWS_COEF molecular weights were calculated from SEQRES records and the highest probability prediction was selected.

Despite the different architectures, the machine learning models are found to give largely similar performance. Given the non-uniform distribution of solvent values found in the PDB (Fig. 3), it is useful to break the performance down for different solvent ranges. Fig. 6 shows the RMSE of each model split into 10% intervals. Interestingly, for structures with solvent content around 40%, MATTHEWS_COEF performs as well as the best machine learning model (U-Solv). Outside of this range, however, the performance of the U-Solv model remains significantly more consistent, giving more confidence to the user that the model can be trusted for structures towards the tail ends of the solvent content distribution.

**Figure 6.**
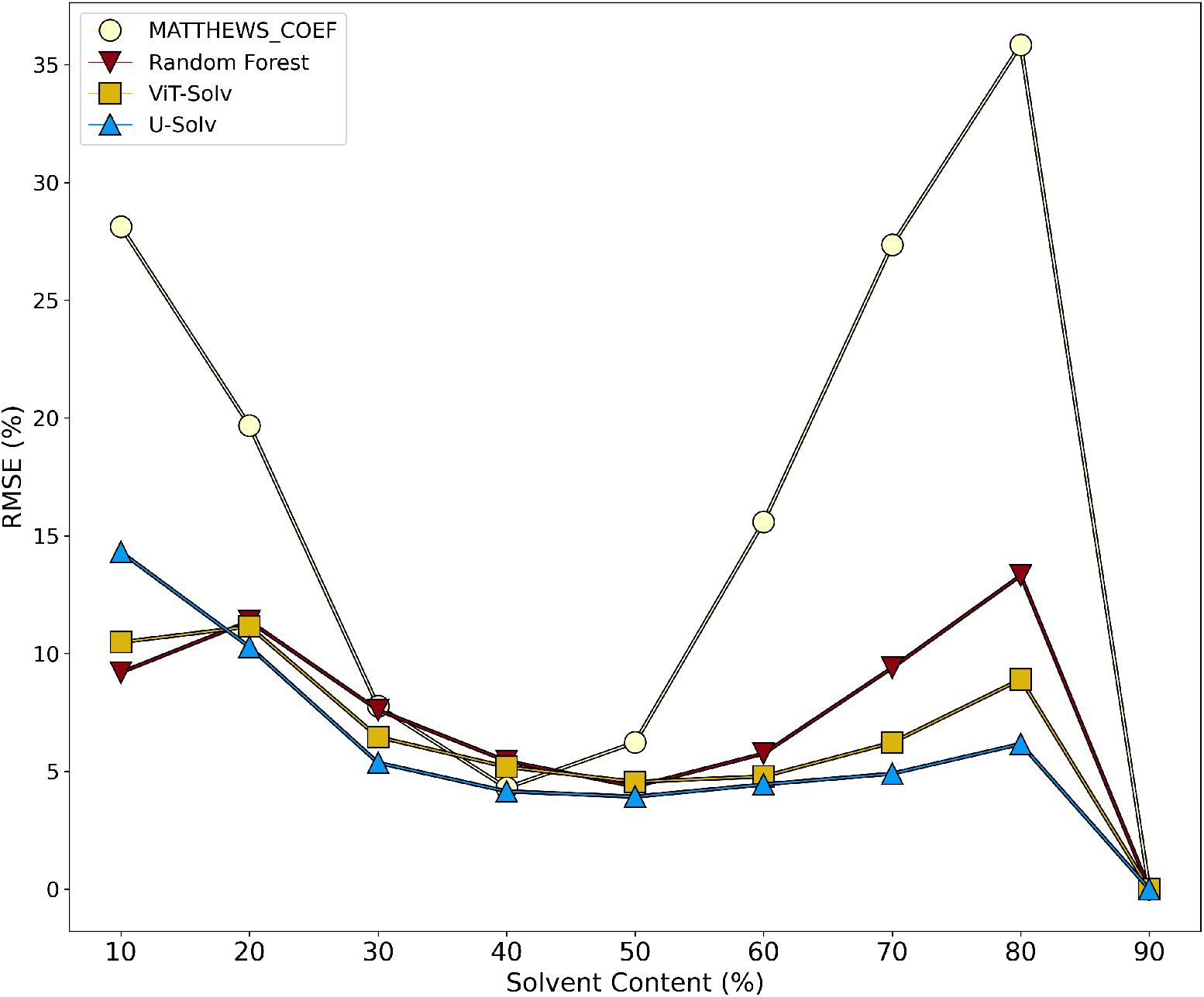
The RMSE of each of the four models split into 10% intervals of the reported solvent content.

Using the solvent content predictions from the best machine learning model (U-Solv), these values can be used within MATTHEWS_COEF to select the most likely number of molecules in the asymmetric unit. Rather than select the number with the highest probability in MATTHEWS_COEF, the number closest to the U-Solv prediction can be selected. This is shown in Fig. 7. For the test set, MATTHEWS_COEF is found to predict 22.7% of structures incorrectly, whereas with the U-Solv model this is reduced to 8.4% (Fig. 7a). As with MATTHEWS_COEF, the trend in poorer performance with increasing number of molecules is still observed with the U-Solv model, but the slope is reduced (Fig. 7b).

**Figure 7.**
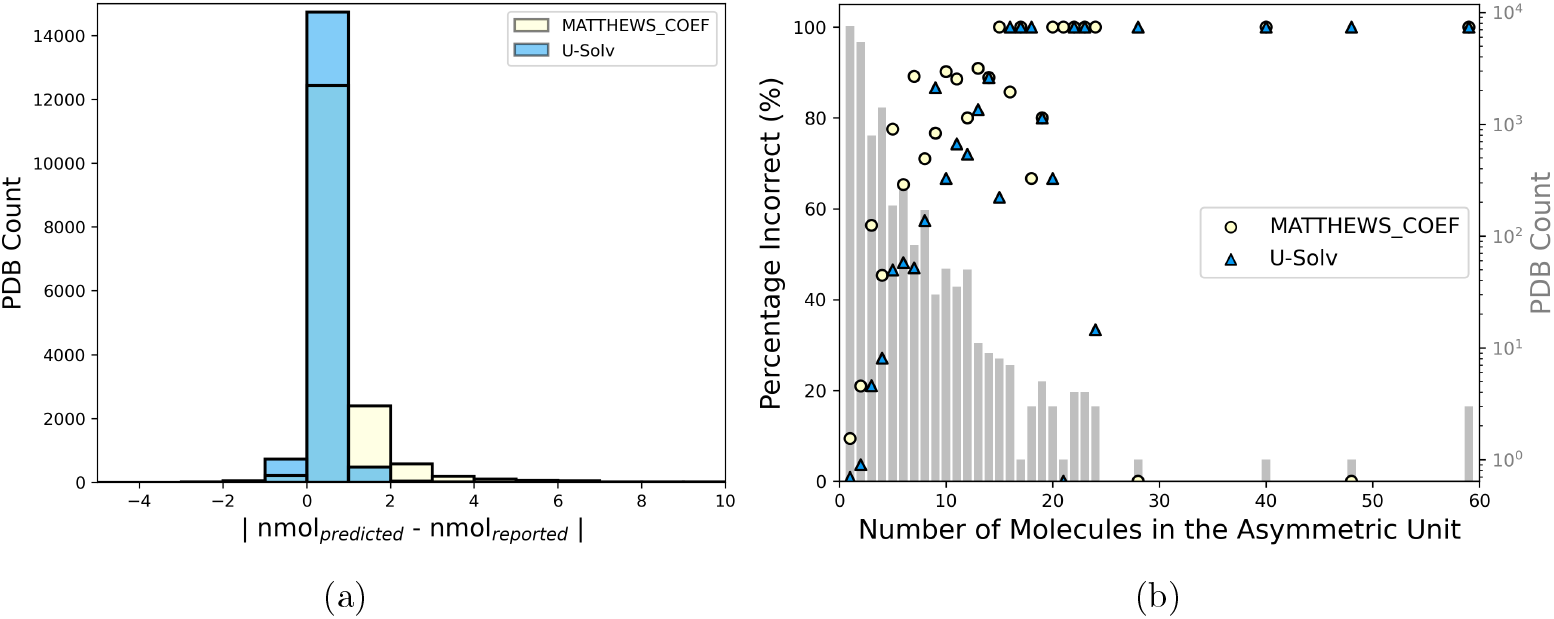
The number of molecules predicted using MATTHEWS_COEF (cream) and MATTHEWS_COEF with solvent content predictions obtained from the U-Solv model (blue) compared to reported values for the test set. (a) A histogram of the differences between predicted and reported values. (b) How the prediction error changes with number of molecules in the asymmetric unit. For MATTHEWS_COEF predictions, molecular weights were calculated from SEQRES records and the highest probability prediction was selected. For the U-Solv results, the MATTHEWS_COEF prediction closest to the U-Solv prediction was selected.

Finally, the uncertainty in the prediction indicated by the RMSE values can be further probed by breaking down prediction error by resolution. Fig. 8 shows the performance of the U-Solv model across different resolution ranges. The trends in increasing prediction error with reduced resolution, and the increasing error at lower and higher solvent content appear instructive, as they seem consistent enough to inform the user of the potential solvent content range to explore, given their dataset.

**Figure 8.**
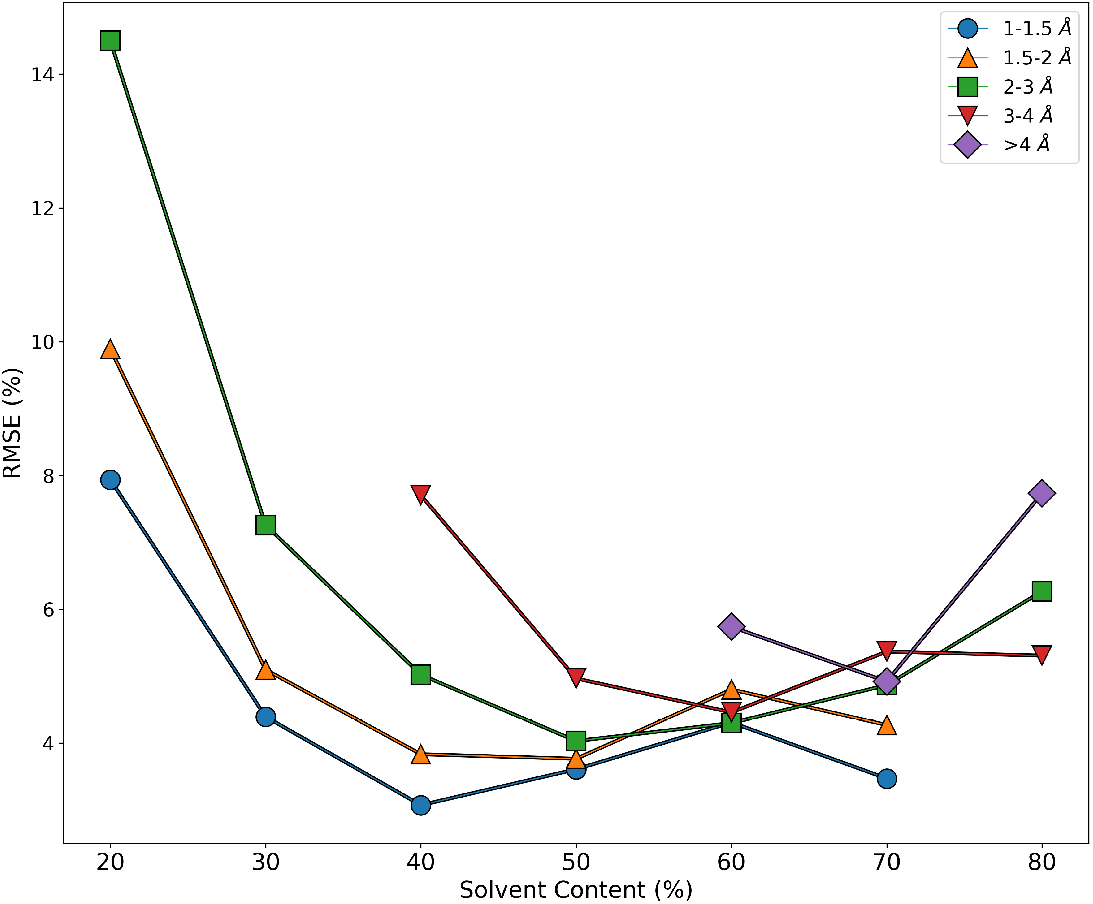
RMSEs of the U-Solv model broken down in terms of solvent content and resolution.

### 3.1 Energy Use

A key motivation for this work was to improve efficiency in the structure solution pipeline, including the energy consumption and hence carbon footprint of the data processing. Although the neural network models outperform the Random Forest model, the differences in training times and resource use are significant. Both neural network models were trained using two NVIDIA RTX A6000 GPUs, with basic data management running on an AMD EPYC 9654P processor. As the training times were significant for both network models, it was not deemed feasible to restrict training to times of low carbon intensity on the national grid. Although the ViT-Solv model ran for nearly twice the epochs as the U-Solv model, individual epoch training times were lower, resulting in only a minor increase in overall training time. For the Random Forest model, all descriptor generation and training (including TPOT calculations) were run on an Intel I7-10875H processor. Although the training time was only a few minutes for the Random Forest model, the TPOT pipeline was run for 24 hours to obtain the optimum machine learning model, based on the training data. Despite this, the carbon emissions of the Random Forest model are observed to be negligible compared to the two network models, with ViT-Solv and U-Solv having carbon emissions higher by a factor of around 50. Although this could not be ascertained beforehand, the improvement in going from the Random Forest model to U-Solv is quite minor (although this becomes greater at higher solvent content (Fig. 6)). This highlights how traditional shallow machine learning models should still be investigated first for a problem, as the result may be sufficient to avoid more expensive approaches.

## 4 Conclusion and Future Work

This work has demonstrated that machine learning models can be readily designed to exploit information from Patterson Maps for predicting solvent content. The best model in this work more than halves the error of current methods, which should lead to significantly more efficient data processing pipelines. As a result, U-Solv is now deployed as part of CCP4 Cloud [38] as the default method. Future development of the method will incorporate nucleic acid datasets, and investigate how assumptions about molecular content could be incorporated as additional parameters. All code used to generate this work is available at www.github.com/ccp4/predict_solvent_content.

The benefits of models like U-Solv can likely be extended beyond predicting a single property of interest. The final connected layer of U-Solv, for example, gives the Patterson Map reduced to a vector component. Outputting this for all structures, this vector space likely contains information that can be exploited for clustering purposes. An indication of this is given in Fig. 9, where the vector output of each structure is further projected down to two dimensions using the UMAP dimensionality reduction method [39].

**Figure 9.**
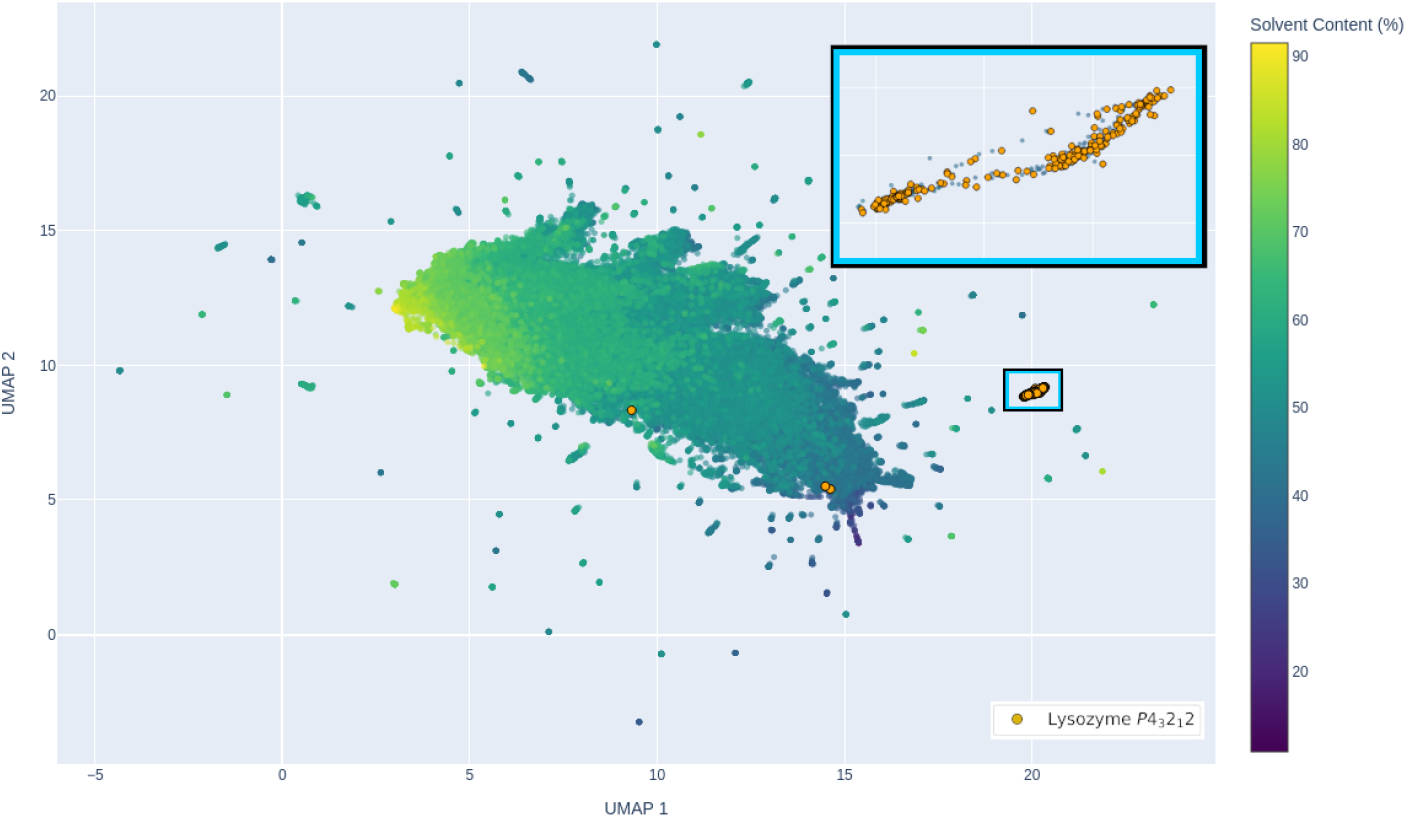
An example projection of the vector space obtained from U-Solv, projected using the UMAP method [39]. Orange markers show Lysozyme *P* 4_3_2_1_2 structures from the PDB. The cluster highlighted in the blue and black box is enlarged in the top-right corner for clarity.

**Figure 10.**
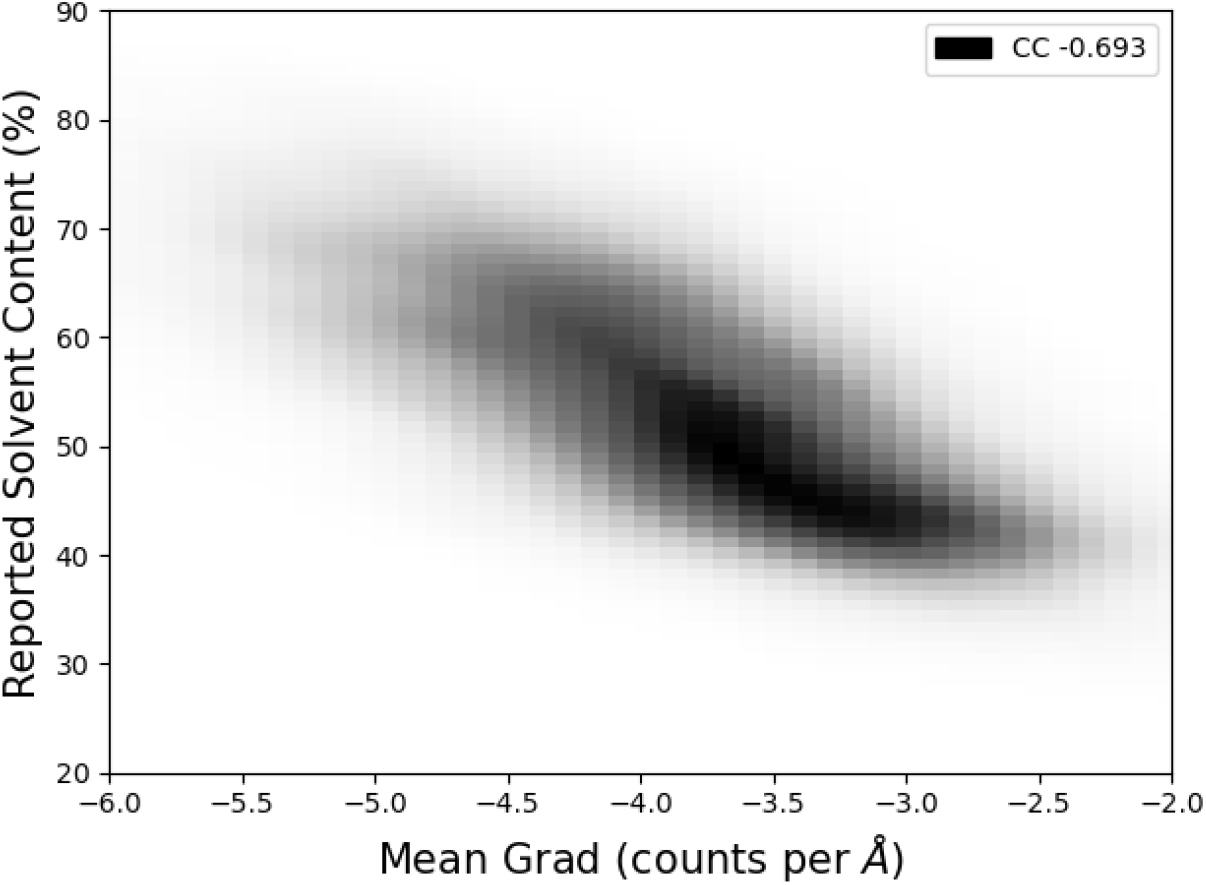
The mean of magnitude of the gradient of the Patterson map, calculated using the Sobel kernel [28] as a 3D convolution.

**Figure 11.**
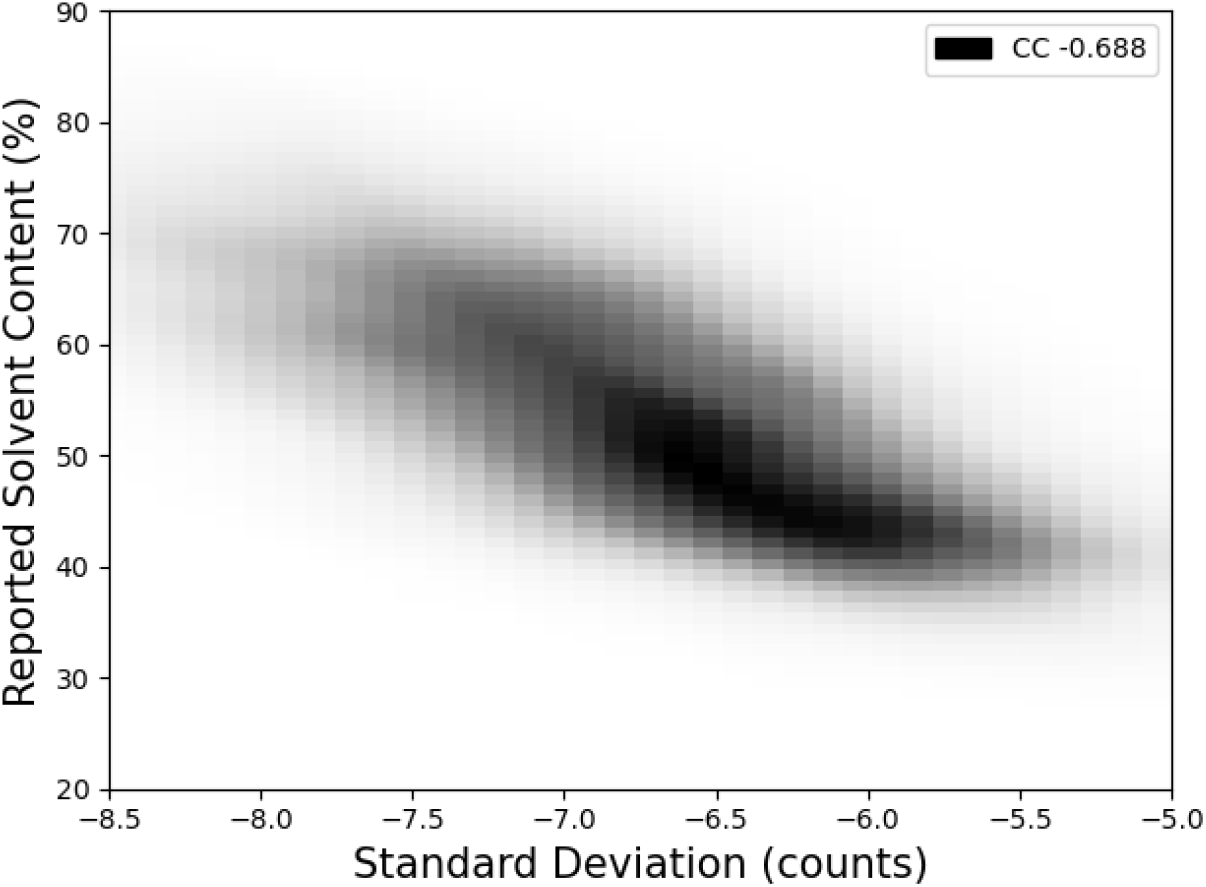
The standard deviation of the Patterson map.

**Figure 12.**
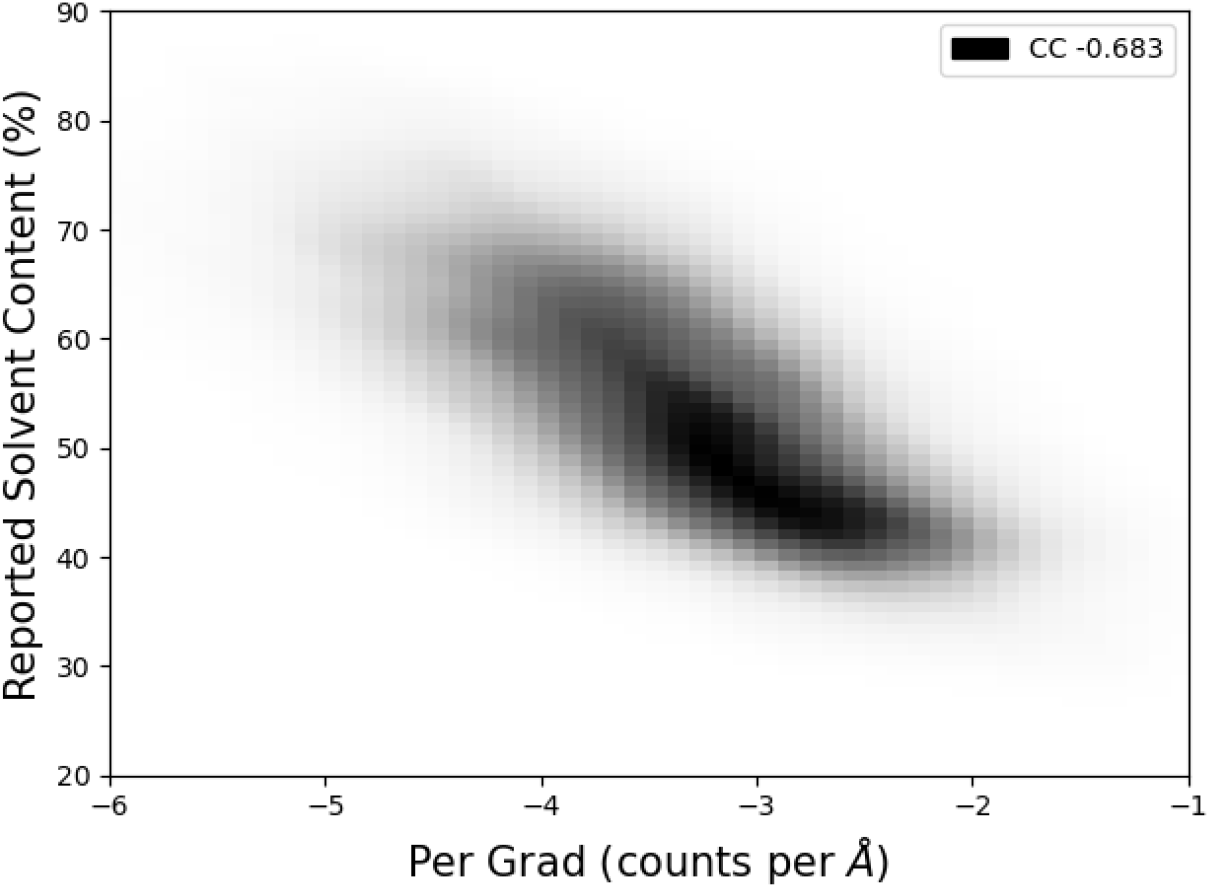
The 90th percentile of the mean of the magnitude of the gradient of the Patterson map, calculated using the Sobel kernel [28] as a 3D convolution. The percentile used was optimised with the training set to maximise correlation with solvent content.

**Figure 13.**
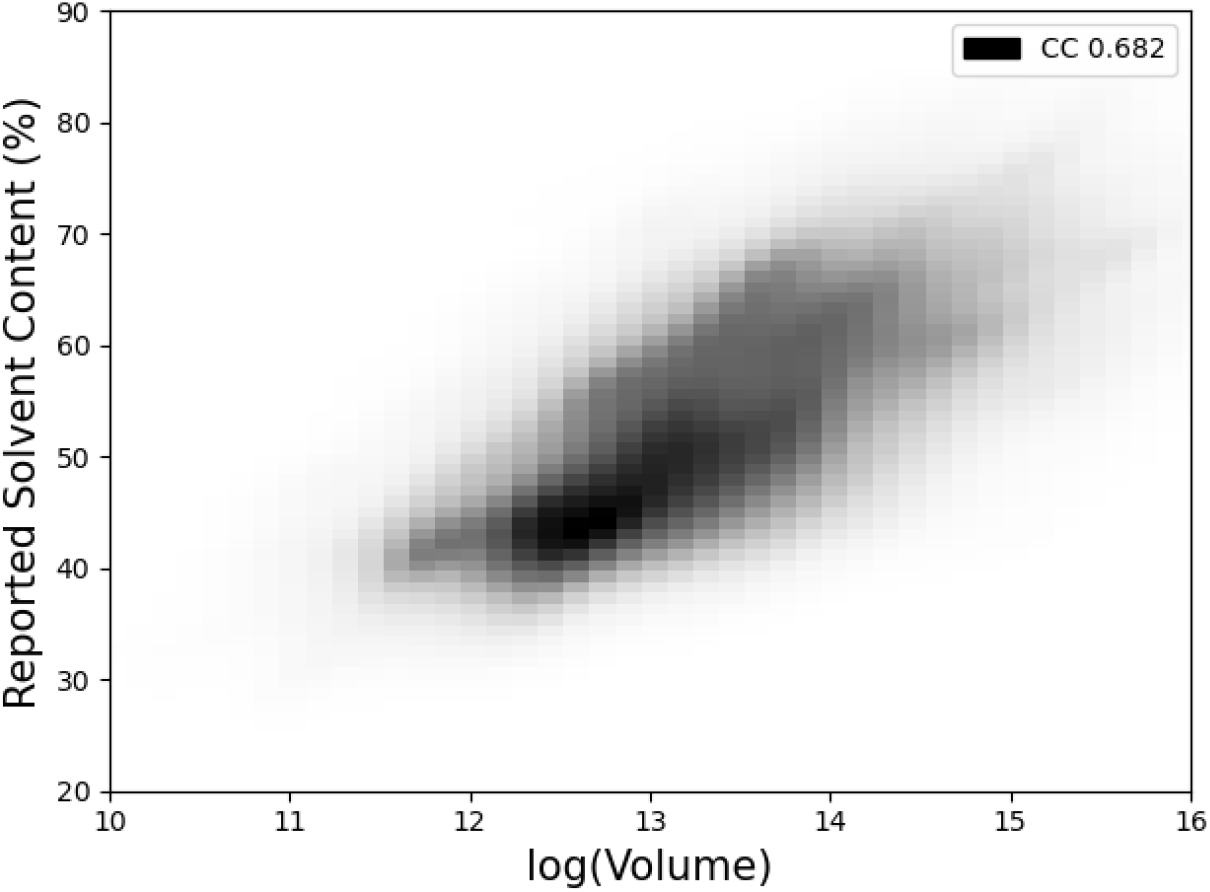
The log of the volume of the unit cell.

**Figure 14.**
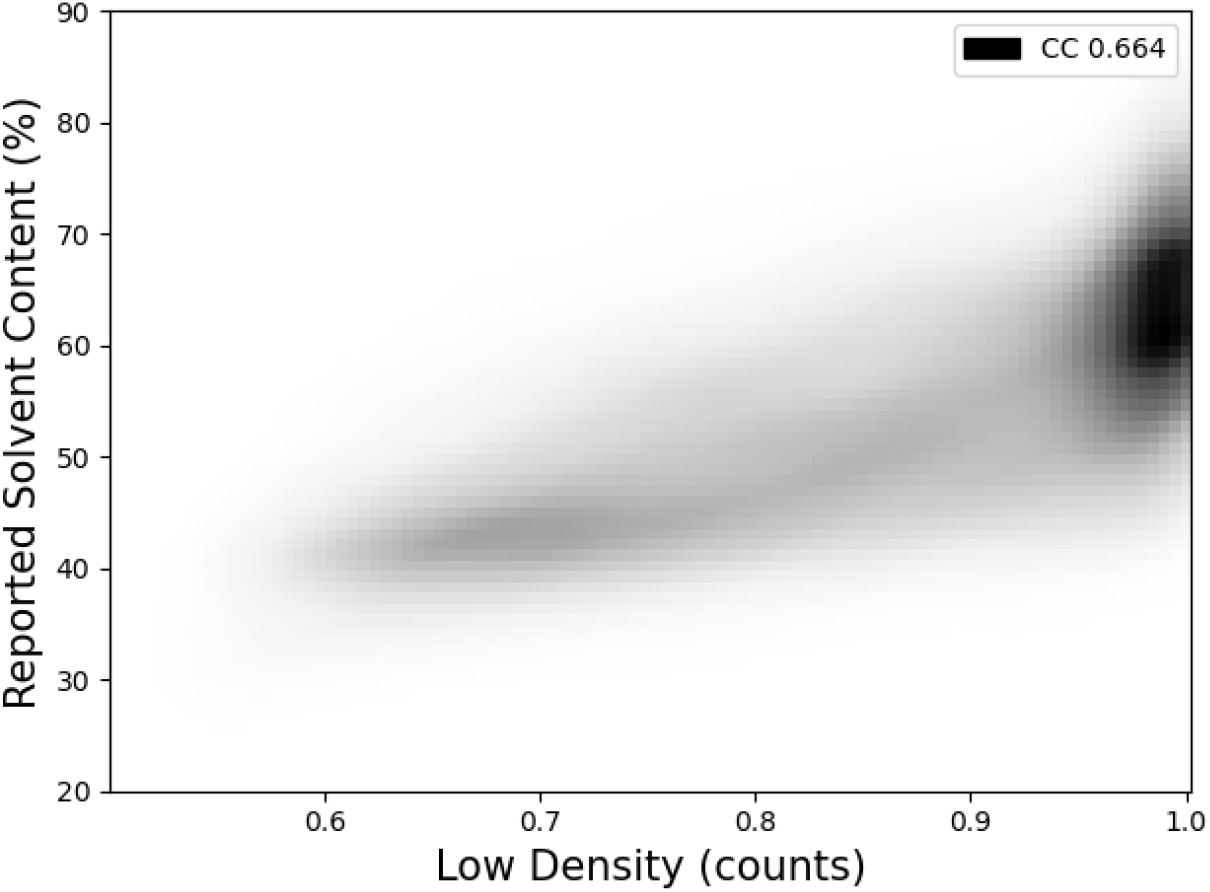
The proportion of values in the Patterson map below a given threshold, normalised with respect to the size of the map, optimised using the training set to maximise correlation with solvent content.

**Figure 15.**
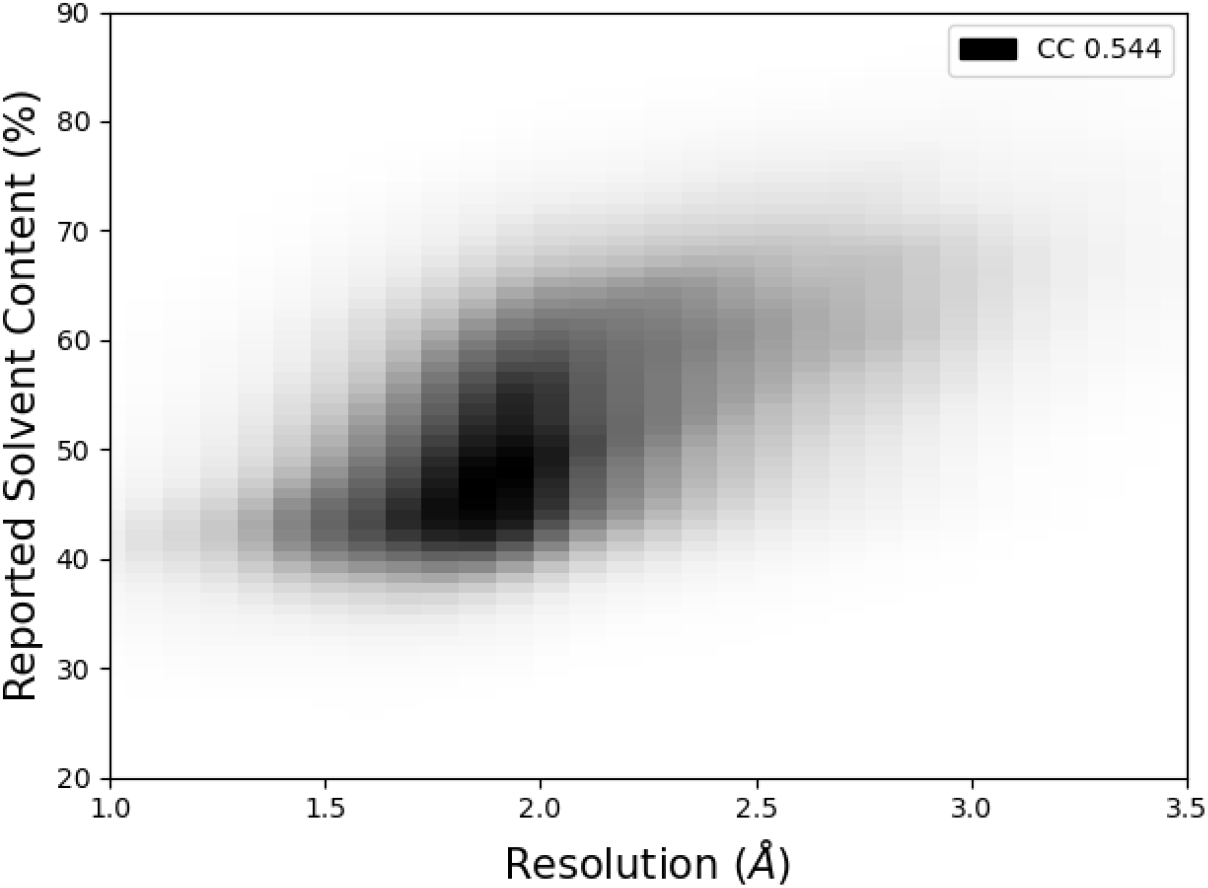

**Figure 16.**
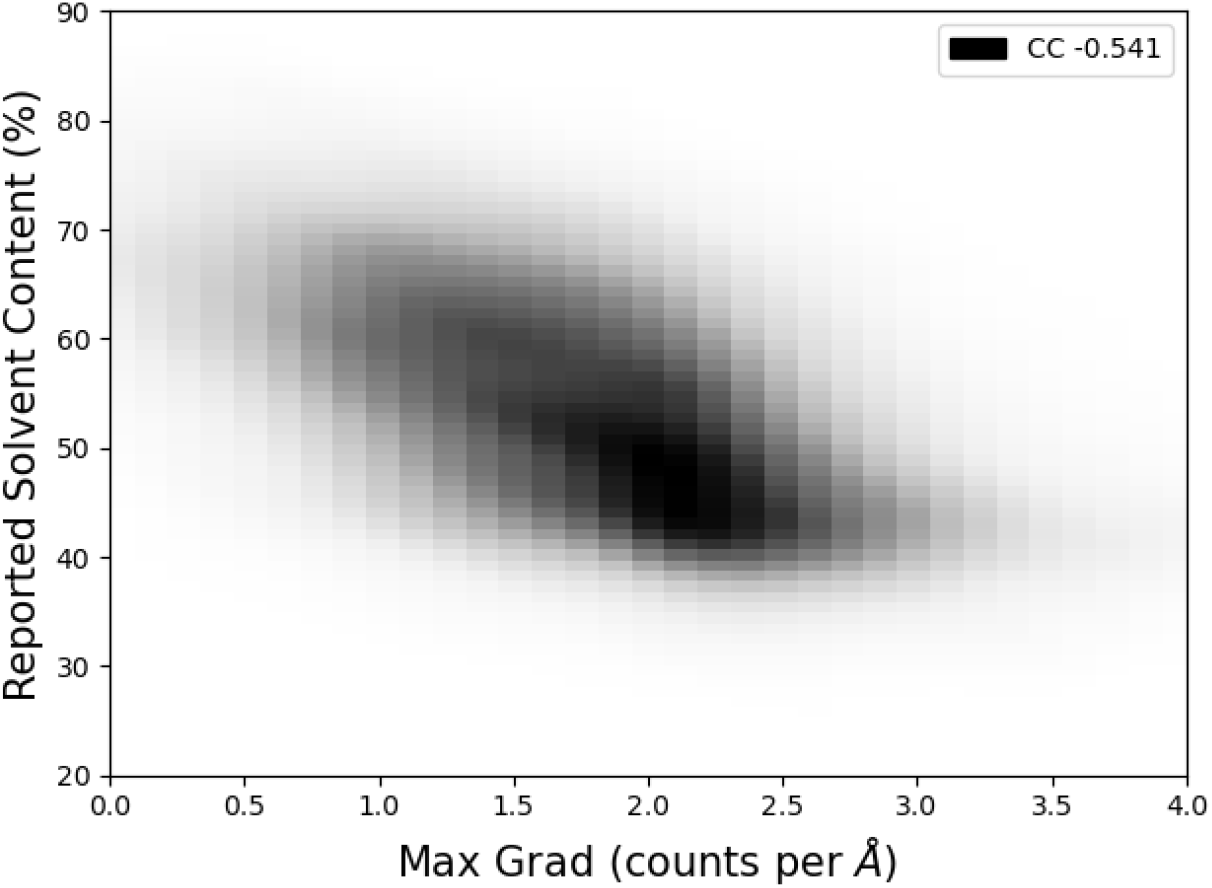
The max of the magnitude of the gradient of the Patterson map, calculated using the Sobel Kernel [28] as a 3D convolution.

**Figure 17.**
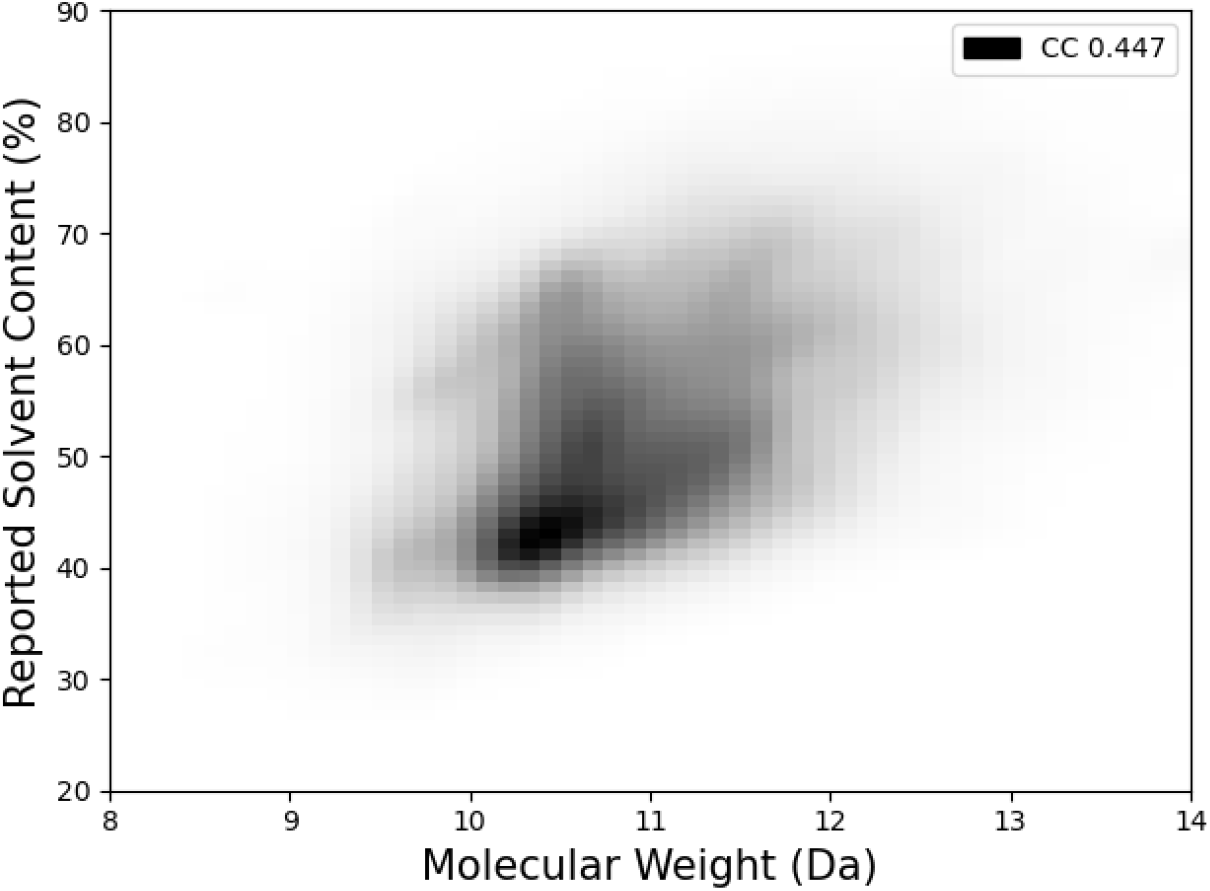

**Figure 18.**
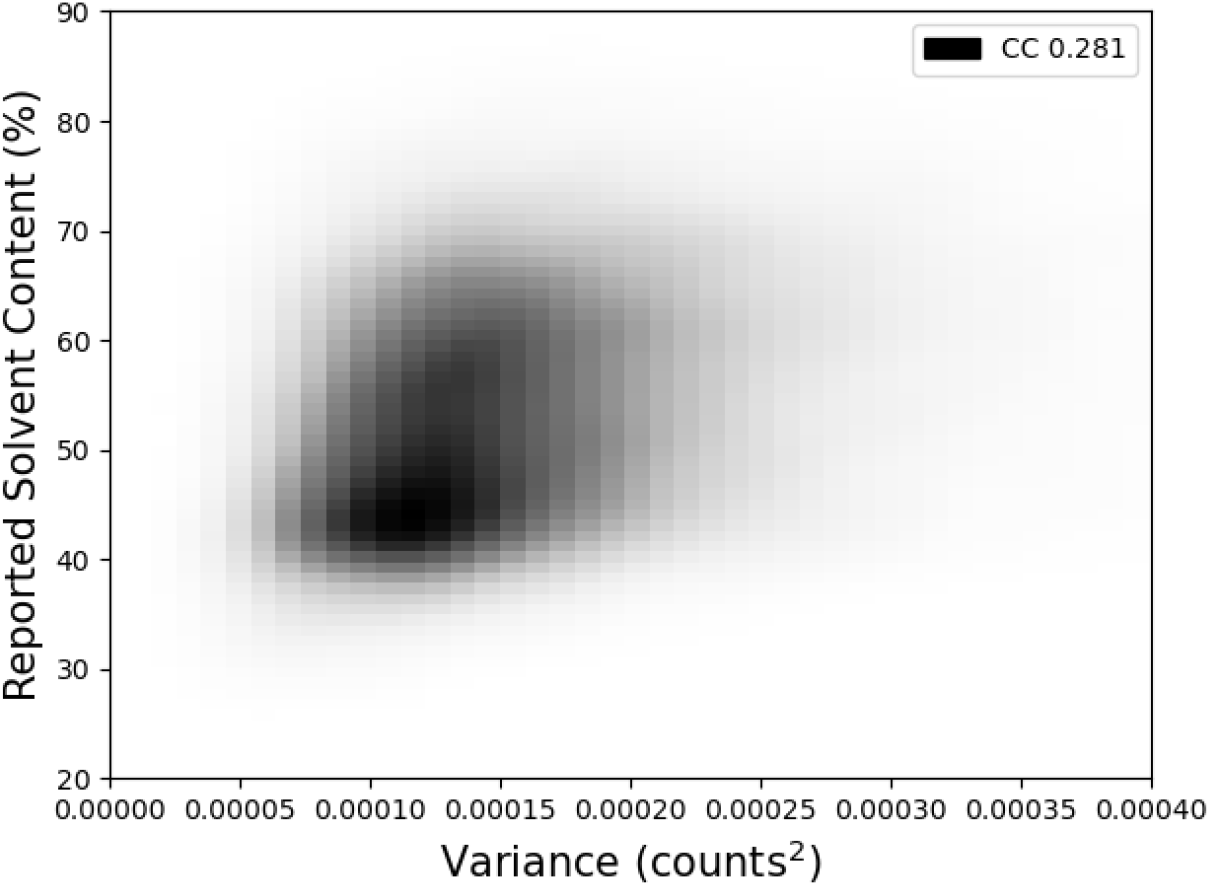
The variance of Patterson map values as a histogram normalised by the total number of bounds, with bin width size optimised using the training set to maximise correlation with solvent content.

**Figure 19.**
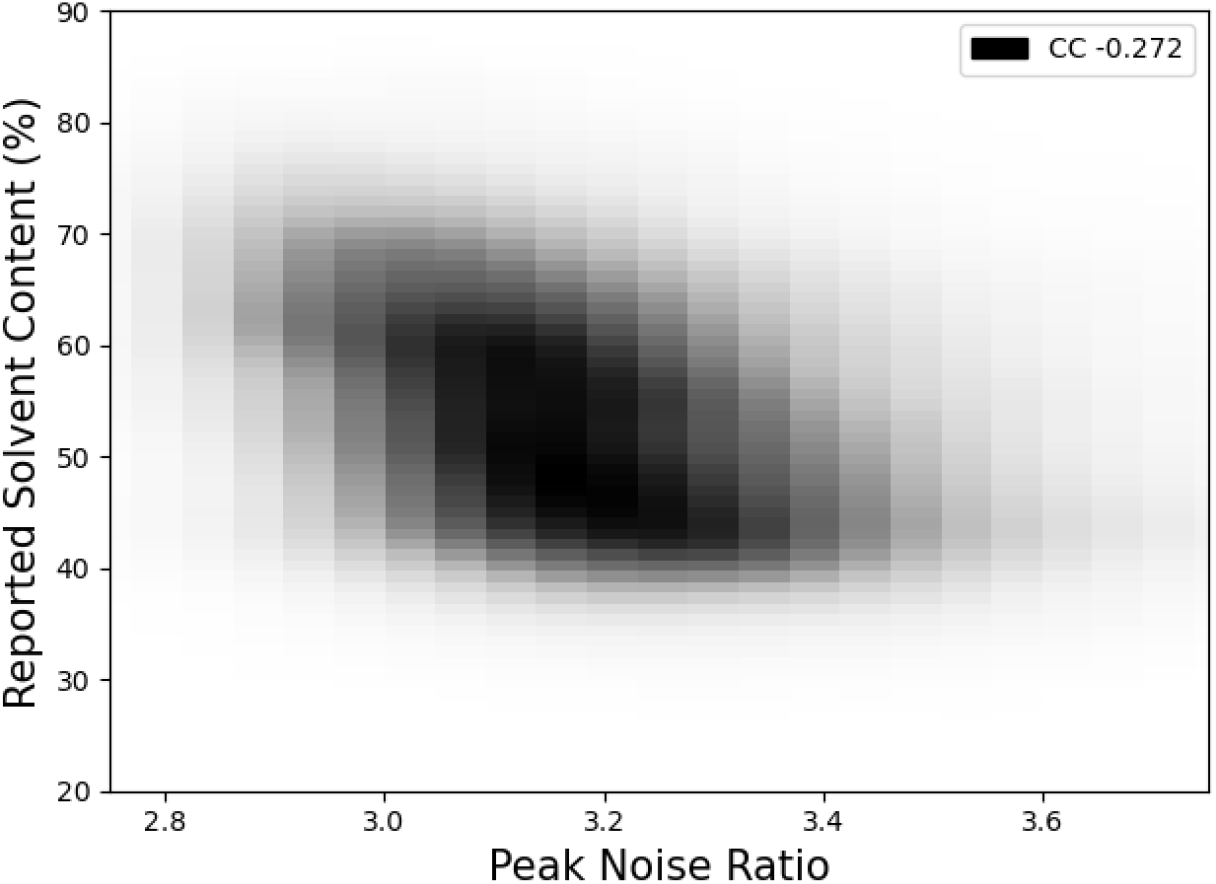
An estimate of the peaks in the Patterson map relative to noise. Peaks were taken as values above *μ* + 2*σ*. The ratio was then calculated as 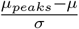

**Figure 20.**
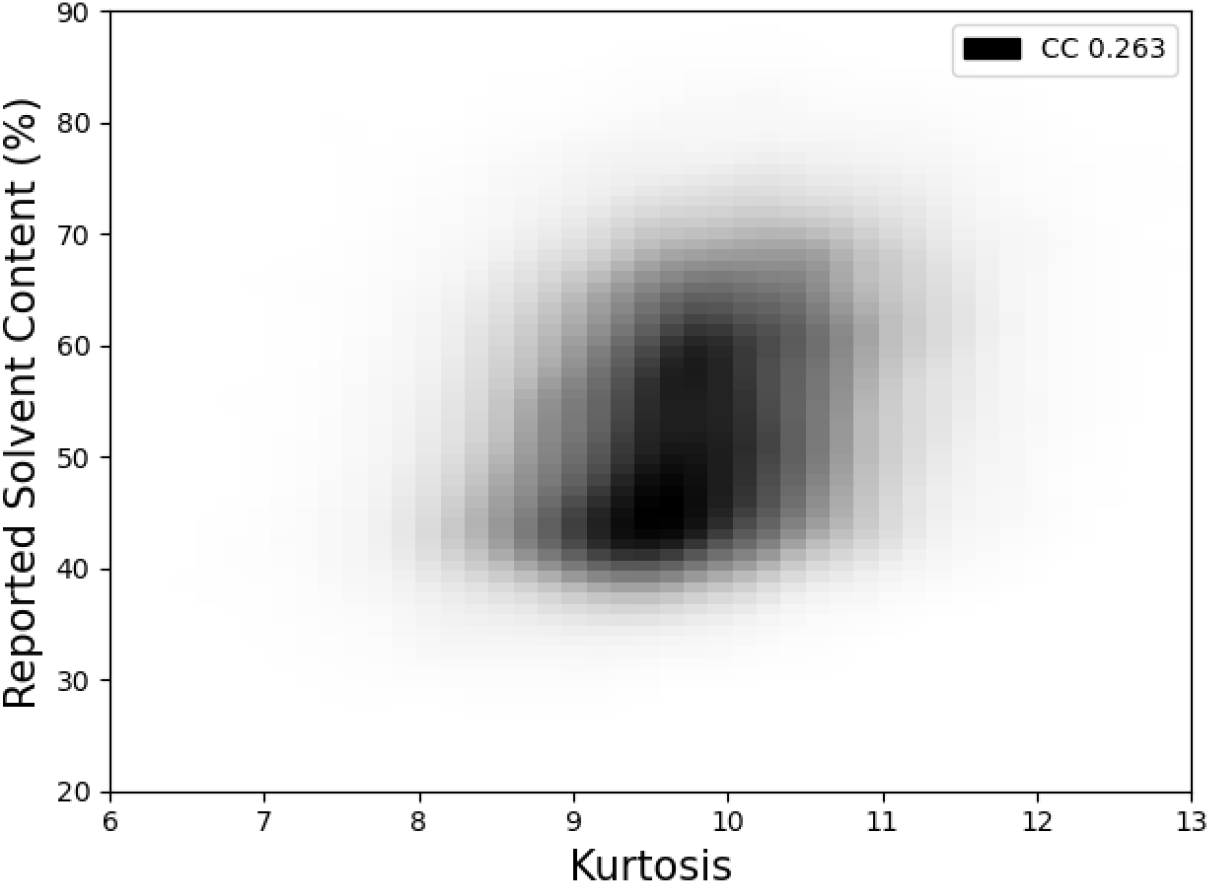
A measure of the distribution of values in the Patterson map defined as 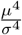, where *μ* is the mean and *σ* is the standard deviation.

**Figure 21.**
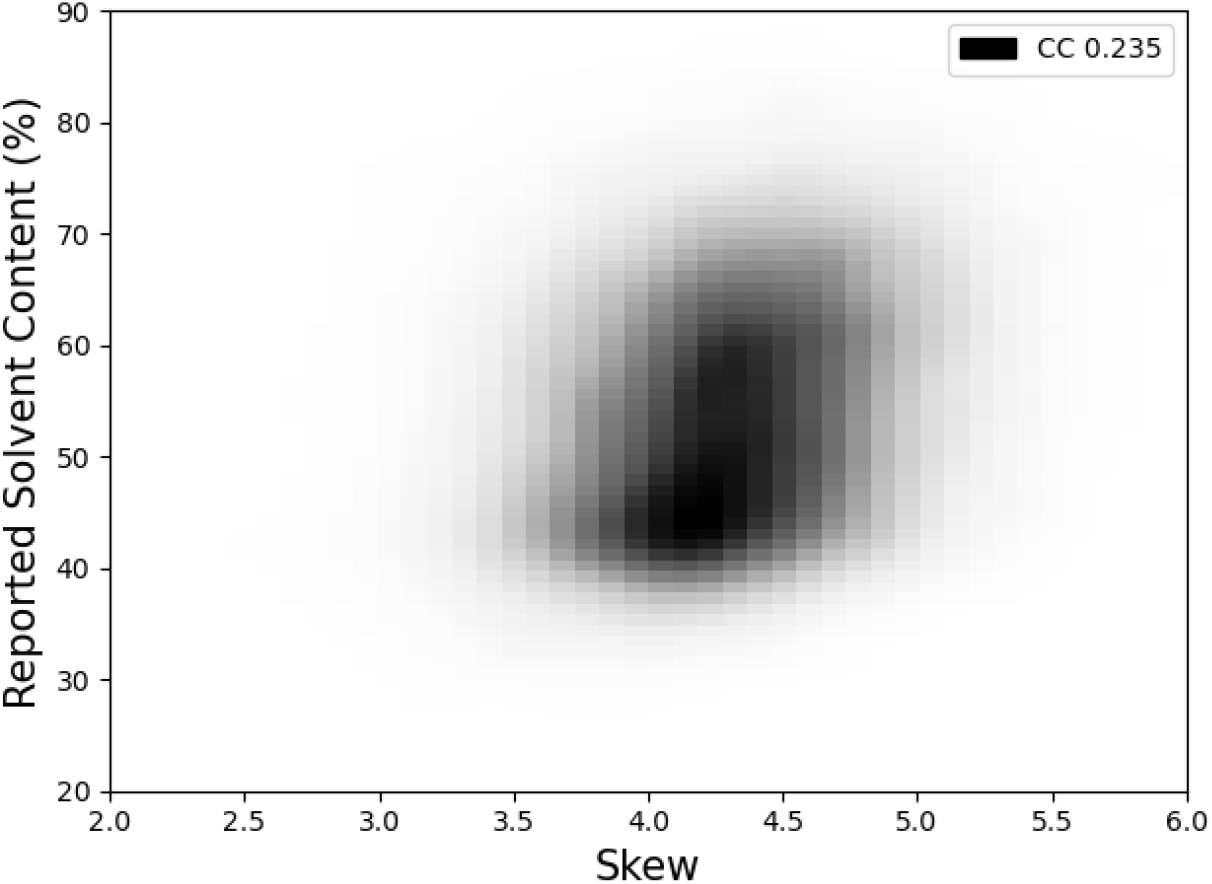
A measure of the skew of values in the Patterson map.

The projection shows clear areas of structure with similar solvent content, as would be expected, along with many clusters. To investigate if this is physically meaningful, structures in the PDB for lysozyme *P* 4_3_2_1_2 were labelled (shown in orange). Of the 946 structures obtained, 246 were in the training set. The majority of these structures are found to cluster together, with three clear outliers. All structures clustering together were found to be structures containing lysozyme from hen eggs. The single isolated structure corresponds to PDB entry 1zv5, which is lysozyme in complex with another molecule. This should significantly affect the unit cell contents and hence the Patterson map and so would not be expected to cluster with the main group of structures. The remaining two outliers correspond to 3ab6 and 3ayq, which were obtained from Meretrix lusoria and have a different amino acid sequence to those in the main cluster. This provides some evidence that these vector spaces could be used for applications such as structure similarity and contaminant detection, and are now the basis of further work.

## Acknowledgments

The authors would like to acknowledge Jools Wills and Ville Uski, as well as CCP4 and the Research Complex at Harwell, for providing computational support for the project. David McDonagh is supported by the Ada Lovelace Centre.

## A Correlation Plots of Manually Selected Features

## B U-Solv Architecture

The UNet implementation used in this study was based on the encoder component of ResidualUNetSE3D as defined in the pytorch-3dunet Github repository [40]. This consists of five layers of residual blocks with an initial input size of (256, 256, 256), downscaling through 16, 32, 64, 128 and 256 feature maps. Each residual block consists of a residual calculated using a 1×1×1 3D convolution, followed by two convolutional layers, where each uses group normalisation followed by an ELU activation function [41]. In the latter layer the residual is added before the ELU function. This output is then passed through a channel/spatial attention module [32], which aims to help emphasise different spatial features and feature map channels via rescaling. After every residual block max pooling is used, such that for a single Patterson map the overall progression in dimensionality reduction is

1. (256, 256, 256)
2. (16, 256, 256, 256)
3. (32, 128, 128, 128)
4. (64, 64, 64, 64)
5. (128, 32, 32, 32)
6. (256, 16, 16, 16)

Global pooling is the used to reduce this to a 256 vector, which is then used in a fully connected layer passed through a sigmoid function to give a final value between zero and one. The code used to generate this model is available at www.github.com/ccp4/predict_solvent_content in usolv.py.

## C ViT-Solv Architecture

The ViT-V-Net implementation used in this study was based on the encoder component of the ViT-V-Net 3D model given in the vit-pytorch repository [34]. Using encoder channels 16, 32, and 32, the initial Patterson map array of size (256, 256, 256) is first passed through two convolutional + pooling layers, downsampling the input to 32 feature maps of size (64, 64, 64). These feature maps are then divided into 512 non-overlapping 3D patches of size (8, 8, 8), each projected into a 252-dimensional embedding. Learnable positional embeddings are added, giving a sequence of shape (512, 252). The embeddings are averaged across the 512 patches to obtain a single 252-dimensional vector, which is passed through a fully connected layer and sigmoid activation to output a value between zero and one. The code used to generate this model is available at www.github.com/ccp4/predict_solvent_content in vitsolv.py

## Notes

https://github.com/ccp4/predict_solvent_content

### Competing Interest Statement

The authors have declared no competing interest.

